# Practice modifies the response to errors during a novel motor sequence learning task

**DOI:** 10.1101/2020.10.09.334169

**Authors:** Dhanush Rachaveti, Rajiv Ranganathan, Varadhan SKM

**Author notes:** **Corresponding Author:** Dr Varadhan SKM Assistant professor Department of Applied Mechanics Indian Institute of Technology, Madras Chennai - 600036 Tamil Nadu.

## Abstract

The occurrence of an error when performing a motor sequence causes an immediate reduction in speed on subsequent trials, which is referred to as post-error slowing. However, understanding how post-error slowing changes with practice has been difficult because it requires extended practice on a novel sequence task. To address this issue, we examined post-error slowing in a novel glove-based typing task that participants performed for 15 consecutive days. Speed and accuracy improved from the early to middle stages of practice, but did not show any further improvements between middle and late stage of practice. However, when we analyzed the response to errors, we found that participants decreased both the magnitude and duration of post-error slowing with practice, even after there were no detectable improvements in overall task performance. These results indicate that learning not only improves overall task performance but also modifies the ability to respond to errors.

## Introduction

Mastering real-world skills such as typing or playing the piano involve a specific type of motor learning termed sequence learning^1, 2, 3, 4^. From the viewpoint of task performance, motor sequence learning has been extensively characterized – learning results in overall improvements in speed and accuracy^5, 6, 7, 8, 9^. The underlying neural changes associated with such learning have also been well documented both in typical controls and in individuals with motor disorders^10, 11^.

Although these overall improvements in speed and accuracy are well described, the question of how participants respond to errors on shorter time scales is less understood. As suggested by Crump and Logan (2013)^12^, errors may serve two distinct roles – (i) a prevention role, in which participants learn from errors to prevent future errors, and (ii) a correction role in which participants simply respond in a way to fix the error that was made. A critical distinction between these two roles is based on how participants respond following an error on shorter time scales – the prevention role is characterized by ‘post-error slowing’ (i.e. an increase in movement time following an error)^13, 14, 15, 16, 17, 18^.

A critical question is whether this post-error slowing is modified with learning. Logan and Crump (2013) suggested that practice would result in a decrease of post-error slowing, possibly because there is not much to learn from an error at higher skill levels. However, to date, this evidence has mostly been cross-sectional^19, 20, 21, 22, 23, 24, 25, 26^. There are two limitations of such cross-sectional designs: (i) the ability to make causal inferences about the role of practice is limited, and (ii) measurements of post-error slowing are confounded with changes in the level of absolute task performance - i.e. because novices are slower than experts, comparing the ‘amount’ of slowing can be a challenge because of the differences in baseline performance.

To address these limtations, in the current study, we examined post-error slowing in a longitudinal design using a novel sequence learning task. Participants practiced a glove-based typing task with relatively high complexity (over 250 5-letter words) for an extended period (15 days). This unique experimental design allowed us to (i) examine causal effects of practice on post-error slowing, and (ii) minimize confounds of task performance by examining changes in post-error slowing after task performance has reached a relative ‘plateau’. Based on the prior literature on post-error slowing, we tested the hypothesis that post-error slowing decreases with practice^21^.

## Methods

### Participants

Eight healthy, right-hand dominant participants (6 males & 2 females, M ± SD: Age: 26.8 ± 2.6 yrs., Height: 163.5 ± 6.0 cm, Weight: 66.8 ± 10.5 kg) volunteered for the study. Participants had no history of any neuro-motor disorder or trauma to the hand or fingers and were naïve to the purpose of the experiment. Handedness was determined using Edinburgh handedness inventory^27^ and all participants had a handedness score above 90% (score of 90 and above indicates that participants were right hand dominant). All participants provided written informed consent before participating in the experiment. The institutional ethics committee of the Indian Institute of Technology Madras approved all the procedures needed to conduct this study (Approval number: IEC/2016/02/VSK-12/22).

### Experimental Setup

Our experimental system was a glove-based typing device (Figure 1a). This system consisted of a glove with conductive key patches placed approximately at the centre of each segment in the index, middle, ring and little fingers (3 segments * 4 fingers = 12 keys) and one at the distal end of the thumb. Among these 13 keys, the one on the thumb was used as a switch while the 12 on the other fingers were assigned with nine specific letters, space, backspace and caps lock. These keys were connected to a microcontroller (Teensy 2.0++) using conductive thread, metallic buttons and cables.

**Figure 1:**
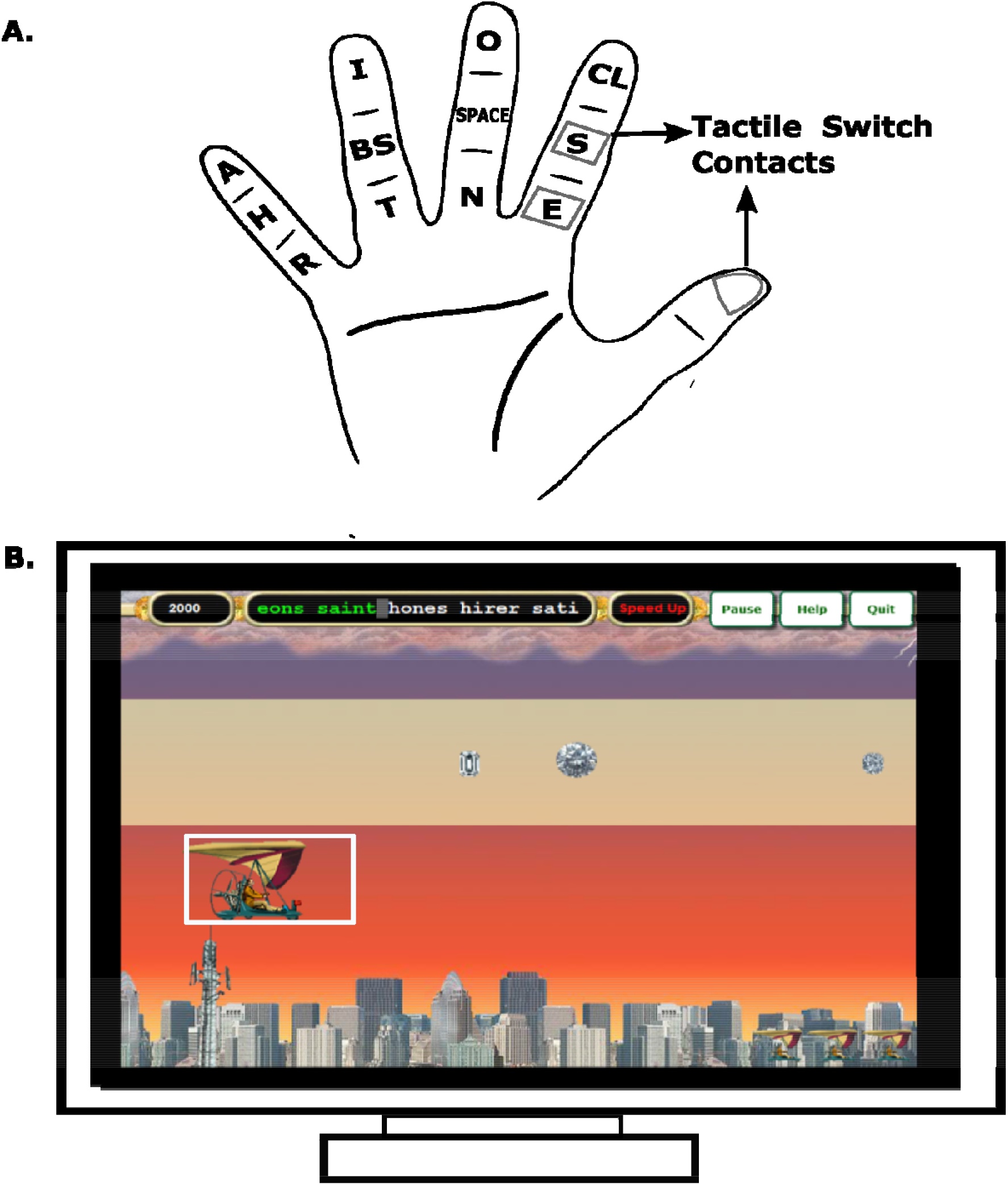
Schematic of the experimental setup. **(A) Glove based typing device**. Participants wore gloves (showed with dotted lines) such that the tactile switches on the glove faced towards the participants. Key patches (shown as squares around the alphabets) were sewn on each finger segment on the gloves. SP, CL, BS denotes “SPACE”, “Caps Lock”, “Back Space”. Conductive threads sewn on the glove were used to connect the key patches to a button connector, which was used to interface with the microcontroller. When a key patch was touched with the thumb, a custom-written code in the microcontroller converted the touch into text. This was shown on the computer monitor **. B. Practice interface:** Words were typed in a game environment with words moving from from right to left as participants typed the words. The objective of the game was to type the words as fast and accurately as possible so that the glider (highlighted in white box) moved from left to right towards the destination. There was also a speed and accuracy constraint - the glider lost altitude and eventually crashed if participants did not maintain a particular speed or accuracy. The words to be typed were shown to the participants in the top panel of the game. The correctly completed keys and words were highlighted in green, and the words yet to be typed were shown in white. At the end of each word participants typed “SPACE” to move to the next word. The Caps Lock and Back Space keys were never used in the current experiment.

To type a particular key, participants had to touch the corresponding key patch on a finger with the thumb, which then closed the electrical circuit. A customized program in the microcontroller detected this event, and the program then sent the ASCII code assigned to that specific key to the computer through a USB port. For example, when the participant touched the middle phalanx of the index finger, the letter ‘S’ was typed on the computer screen, (Figure 1a). Gloves were custom-made to suit the hand dimensions for each participant. The text from the glove was processed by a customized LabVIEW based program at 1000 Hz.

### Task

The goal of the participants was to type a set of words as fast and as accurately as possible. Rather than let participants choose their speed (which might not reflect their actual maximum speed), we selected a ‘game interface’ where we could set the typing speed. This allowed us to probe the maximum speed of the participants more closely. This interface is described below.

### Training words

Words used in the experiment were 5-letter words picked from a custom dictionary. This dictionary comprised of 281 words each made up from the nine most frequently used letters - e, s, o, n, i, t, a, r, h^28^. These letters were mapped to keys on the glove to form a ‘key-map’(Figure 1b). For all participants, the same key-map was used on all days and blocks

### Game Interface

The words typed by participants were displayed in a game interface (Diamond glider game in Typing Instructor^®^ Platinum 21, Individual software, CA, USA) (Figure 1c). The objective of the game was to type words quickly and accurately to move a glider from the starting point to destination without crashing. The glider moved towards the destination as the participants typed. Words to be typed appeared on the right and moved left on the screen as the participant typed them. If a correct letter was typed, that letter was highlighted in green, and the cursor moved to the next letter. If a wrong letter was typed, that letter was highlighted in red, and the cursor stayed on the same letter, until the correct letter was typed. An audible beep tone was played when an error occurred. In addition to the words, participants had to type the SPACE key in between words (For more details see our data paper^28^).

### Protocol

Participants practiced the experimental task for 15 consecutive days (including weekends) and data were collected on all days of practice. Each day/session was divided into 12 blocks of 2 mins each with 30 seconds interval between blocks. Words could repeat within a block but not between blocks, and words on a given block remained same across all days (i.e.. the m^th^ Block was composed of the same set of words on all days but the order of word presentation may change between days; the n^th^ block always had a set of words different from the m^th^ block, when m≠n). All blocks had 23 words with the exception of the 12^th^ block which had 28 words.

## Data Analysis

### Movement Time

Movement time (MT) was defined as the time taken to reach/press a particular letter after the release of the previously typed letter, which is computed as the difference between keypress time of the specific letter and the key release time of the previously typed letter. This value was averaged across all blocks in a given day of practice.

### Errors

Errors were defined as the ratio of the number of letters mistyped to the total letters typed in a block. This value was averaged across all blocks in a given day of practice.

### Post-error slowing

Post-error slowing was assessed using two measures: (i) the magnitude, which refers to the increase in MT after an error. and (ii) the duration, which refers to the time taken (measured in keystrokes) for the MT to recover to pre-error levels. To separate “pre-error” vs. “post-error” segments, we first traversed to every error in a block and separated the MT values into segments before (pre) and after (post) the onset of an error. For each error, we then determined a ‘recovery point’ by examining the point where the post-error MT was equal or less than the pre-error MT.

### Magnitude

The magnitude of post-error slowing was computed as the absolute difference between the average of MT values before and after an error. For the ‘pre-error’ segments, the average of MT values prior to an error was considered until the recovery point of the previous error (or to the first keystroke if this was the first error). For the ‘post-error’ segments, the average of MT values after an error was taken until the recovery point of the current error (or to the last keystroke if this was the last error). Because MT also decreases with practice, we computed the magnitude of post-error slowing as a ‘relative change’ by normalizing the change in MT (the absolute difference between mean values of MT before and after an error) to the mean values of MT just before an error. Thus, if the magnitude of the post-error slowing reduces, it indicates a smaller decrease in MT after an error.

### Duration

The duration of post-error slowing was computed as the number of keystrokes it took for the MT value to become less than or equal to pre-error MT values. Similar to the magnitude computation, the duration was defined based on the recovery point. Thus, a reduction in the duration of post-error slowing indicates that MT took less time to recover to pre-error values.

## Statistical Analysis

Data from the outcome variables were organized into three stages – the Early-stage consisted of Days 1, 2, 3; the Middle stage included Days 7, 8, 9, and the Late-stage consisted of Days 13, 14, 15. One-way repeated-measures ANOVA with three levels was used for statistical analyses on all the outcome variables with Practice stage as a factor (3 levels – Early, Middle, Late) and participant as a random factor. Corrections for sphericity were performed using the Huynh-Feldt criterion wherever appropriate. Significant effects were further analyzed using Tukey’s posthoc test. Effect sizes are reported using partial eta-squared values (η_p_^2^).

## Results

Overall, participants showed improvements in task performance as they practiced the typing task. This performance improvement was seen as an overall decrease in the movement times (Figure 2a) and errors (Figure 2b).

**Figure 2:**
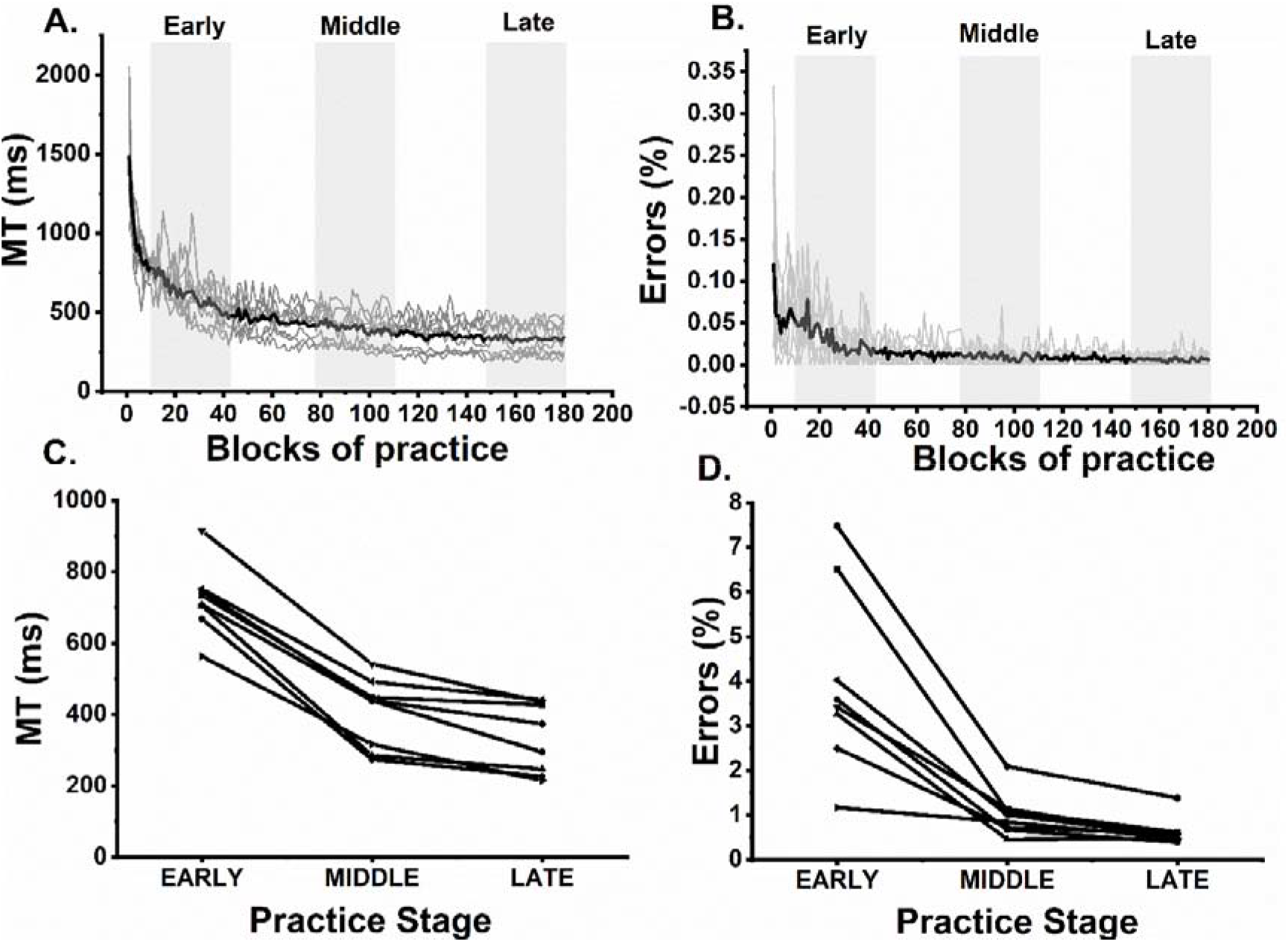
Changes in task performance with practice, (A) Movement time and (B) Error percent are shown as a function of practice for all the participants. The black line indicates the mean across the participants while the grey lines indicate data from individual participants. Each day of practice consisted of 12 blocks. The first three days (36 blocks) were considered as the early stage of practice, days 7 to 9 were considered as a middle stage of practice, while the last three days were considered as a late stage of practice (C) MT as a function of practice stage for individual participants. Movement time (MT) reduced significantly from the early to the middle stage, but there was no significant difference between the middle and late stages of Practice (D) Error percent as a function of practice stage for individual participants. Error percent reduced significantly from early to late stages of practice. Each line represents a single participant. There was a significant improvement from early to the middle, but there was no significant difference between middle and late stages of practice.

### Movement Time

MT reduced with practice (Figure 2c). This observation was supported by one-way repeated measures ANOVA that showed a reduction in MT due to practice stage (F_(1.82, 12.74)_ = 179.30, p <0.001, η_p_^2^=0.96). Post-hoc comparisons showed that MT reduced between early and middle stages (mean MT: Early-724 ms, Middle-404 ms and Late-332 ms. p < 0.001), but there was no significant difference between middle and late practice stages (p = 0.16).

### Errors

Errors reduced with practice (Figure 2d). This observation was supported by a one-way repeated measures ANOVA that showed a reduction of errors due practice stage (F_(1.03, 7.21)_ = 25.47, p<0.001, η_p_^2^=0.83). Similar to the movement time results, post-hoc comparisons showed that errors decreased between early and middle stages (mean errors: Early- 3.9%, Middle- 0.9%, Late- 0. 6%; p < 0.001), but there was no significant difference between middle and late practice stages (p = 0.55).

### Post-error slowing

Post-error slowing (i.e. increase in MT following an error) for a single participant can be seen both in terms of the absolute movement time (Figure 3a) and normalized movement time (Figure 3b). The post-error slowing showed changes both in magnitude and duration with practice.

**Figure 3:**
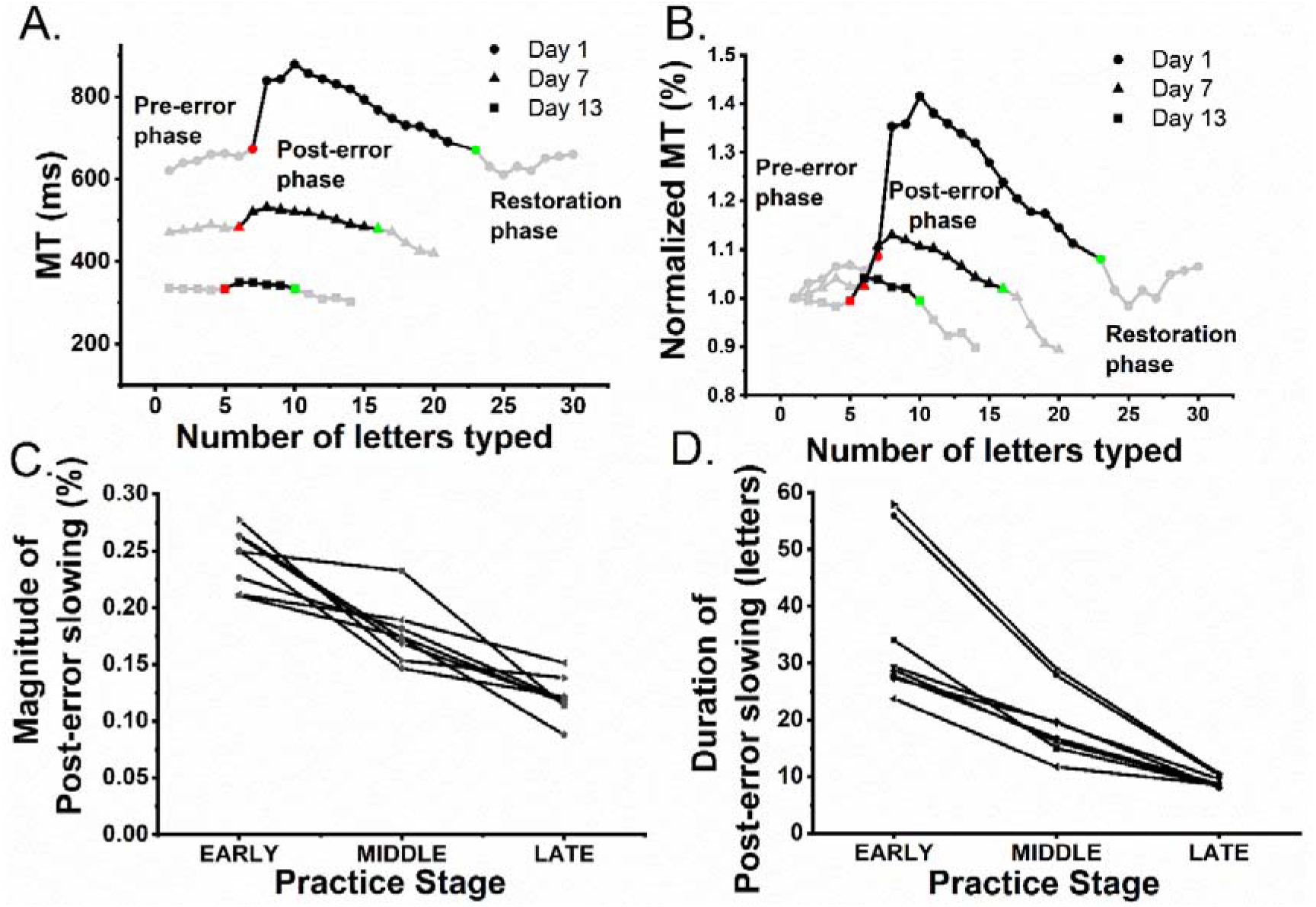
Changes in post-error slowing with practice. **(A)** Raw data of movement time and occurrence of an error from a single participant: Raw MT data plotted on three days (day 1, 7 and 13) and split into three phases (pre-error, post-error, and restoration). The onset of an error is indicated as a solid red symbol, and the onset of the recovery (i.e.) is indicated by the green symbol. **(B)** Normalized raw data of movement time and occurrence of an error from a single participant. The same data in panel A is represented as a ‘normalized’ value by dividing each value by theo pre-error value (first value in the absolute MT graph (section a)) of MT **(C)** Change in the magnitude of post-error slowing across all subjects as a function of practice shows the change in the magnitude of post-error slowing for individual participants. There was a significant reduction (p < 0.001) in the magnitude of post-error slowing, between early, middle and late stages of practice. **(D)** Change in duration of post-error slowing for individual participants. Once again, there was a significant difference (p < 0.001) between early, middle and late stages of practice. Each line in panels **C** and **D** represents an individual participant.

### Magnitude

The magnitude of post-error slowing decreased with practice (Figure 3c). This observation was supported by a one-way repeated measures ANOVA that showed a reduction in magnitude of post-error slowing due to practice stage (F_(2.4, 16.8)_ = 45.088, p < 0.001, η_p_^2^=0.86). Post hoc comparisons showed that early practice stage was different from middle practice stage (mean magnitude: Early-0.24%, Middle-0.17%, Late-0.12%; p < 0.001). However, there was also a significant reduction in magnitude between middle and late practice stages (p < 0.001) even though MTs between the middle and late practice were not significantly different.

### Duration

The duration of post-error slowing also reduced with practice, as shown in Figure 3d. This observation was supported by one-way repeated measures ANOVA that showed a reduction in duration due to practice stage (F_(1.1, 7.7)_ = 36.27, p<0.001, η_p_^2^=0.83). Post hoc comparisons showed that early practice stage was different from both middle and late practice stages (mean duration: E-36 letters, M-19 letters, L-9 letters; p < 0.001). However, once again, there was a significant reduction in duration between middle and late practice stages (p = 0.002) even though MTs between the middle and late practice were not significantly different.

## Discussion

The motivation for the present study was to understand the phenomenon of post-error slowing as a function of practice. We used three key features in our experiment – (a) a longitudinal design to examine the causal effects of practice, (b) a novel glove-based motor sequence learning task with high task complexity to examine the early phase of motor learning, and (c) an extended practice period until task performance reached a relative plateau to minimize the confounds of comparing post-error slowing at different levels of task performance. Based on prior work, we hypothesized that changes in the post-error slowing would decrease with practice, and our results were consistent with this hypothesis.

First, we found expected effects of practice on overall speed and accuracy. Errors reduced with practice in the early to middle (or late) stage of practice. However, there were relatively small differences between the middle and late stages of practice suggesting that errors plateaued approximately around the middle stage of practice. Similarly, MT reduced from early to the late stage of practice and plateaued between middle and late stage of practice. These results on speed and accuracy in performing novel motor sequence learning tasks are consistent with the prior work^5, 6, 7, 9^.

Second, we found effects of practice on post-error slowing, even after task performance had reached a relative plateau. During the early stage of practice, both magnitude and duration of post-error slowing was high, indicating that participants slowed down more often to avoid future errors. However, during later stages of practice, there was a reduction in both the magnitude and duration of post-error slowing. This reduction in the magnitude and duration of post-error slowing indicates that the participants reacted to errors with a smaller increase in speed and regained speed quickly after errors with no major changes in error rates^29, 12, 30, 14, 15^. This reduction in post-error slowing was seen even after MT and errors relatively plateaued during the late stage of practice. These results confirm that the post-error slowing indeed decreased with practice, and was not confounded by underlying changes in task performance.

There are two potential theoretical explanations for why post-error slowing decreases with practice – i.e. why participants become less sensitive to errors. First, as suggested by Logan and Crump (2009)^21^, the decrease in post-error slowing may be a consequence of the fact that there is ‘less to learn’ from an error as participants become more skilled. Responding to errors can be viewed as a ‘credit assignment’ problem^31, 32^ where the nervous system has to estimate the source of these errors. Since errors become less frequent with learning, participants in the late stage of learning may be more likely to attribute an error to the ‘world’ rather than their own ‘bodies’, which would explain why the sudden occurrence of an error does not alter performance dramatically on future trials. This explanation is consistent with work in other domains such as dart throwing, where the response to an error on the next trial is diminished in experts relative to novices^33, 25^.

A second explanation for the reduction in post-error slowing is that continued reliance on ‘error prevention’ could disrupt automaticity of performance. Motor learning has been characterized by a transition from conscious, deliberate performance in the early stages of practice to more automatic performance in the later stages^34^. This transition has been supported in sequence learning by several features such as chunking^35, 36^ and the ability to perform dual tasks^37, 38, 39^. A smaller response to errors could be reflective of the fact that participants tend to maintain automatic performance and do not switch to a conscious mode of control. This could a beneficial strategy because there is evidence that switching back to a conscious control mode could be detrimental to overall task performance^19^.

In conclusion, we showed that motor sequence learning not only involves changes in overall speed and accuracy, but also in how participants respond to errors, both in terms of the magnitude and duration. Our results suggest that theories of sequence learning not only need to describe overall improvements in task performance, but also need to account for these shorter time scale changes in response to errors, and their change with learning. Understanding these responses to errors may provide greater insight into skilled performance and could also be exploited to tailor practice schedules based on skill level.

## Acknowledgements

The Department of Science & Technology, Government of India, supported this work, vide Reference Nos **SR/CSRI/97/2014 & DST/CSRI/2017/87** under Cognitive Science Research Initiative (CSRI) (awarded to Varadhan SKM).

## Author Contribution

**The authors have contributed as mentioned in the below section**

i. Conception or design of the work – Dhanush Rachaveti, Varadhan SKM
ii. Data collection- Dhanush Rachaveti, Varadhan SKM
iii. Data analysis and interpretation- Dhanush Rachaveti, Rajiv Ranganathan, Varadhan SKM
iv. Drafting the article- Dhanush Rachaveti, Rajiv Ranganathan, Varadhan SKM
v. Critical revision of the article- Dhanush Rachaveti, Rajiv Ranganathan, Varadhan SKM
vi. Final approval of the version to be published- Dhanush Rachaveti, Rajiv Ranganathan, Varadhan SKM

## References

1. Clegg, B. A., DiGirolamo, G. J. & Keele, S. W. Sequence learning. Trends in Cognitive Sciences 2, 275–281 (1998).

2. Palmer, C. & Van de Sande, C. Units of knowledge in music performance. Journal of Experimental Psychology: Learning, Memory, and Cognition 19, 457–470 (1993).

3. Palmer, C. & van de Sande, C. Range of planning in music performance. J Exp Psychol Hum Percept Perform 21, 947–962 (1995).

4. Rumelhart, D. E. & Norman, D. A. Simulating a Skilled Typist: A Study of Skilled Cognitive-Motor Performance. Cognitive Science 6, 1–36 (1982).

5. Friedman, J. & Korman, M. Kinematic Strategies Underlying Improvement in the Acquisition of a Sequential Finger Task with Self-Generated vs. Cued Repetition Training. PLoS ONE 7, e52063 (2012).

6. Karni, A. et al. Functional MRI evidence for adult motor cortex plasticity during motor skill learning. Nature 377, 155–158 (1995).

7. Karni, A. et al. The acquisition of skilled motor performance: fast and slow experience-driven changes in primary motor cortex. Proceedings of the National Academy of Sciences 95, 861–868 (1998).

8. Karni, A. The acquisition of perceptual and motor skills: a memory system in the adult human cortex. Cognitive Brain Research 5, 39–48 (1996).

9. Rozanov, S., Keren, O. & Karni, A. The specificity of memory for a highly trained finger movement sequence: Change the ending, change all. Brain Research 1331, 80–87 (2010).

10. Doyon, J. et al. Experience-dependent changes in cerebellar contributions to motor sequence learning. PNAS 99, 1017–1022 (2002).

11. Stickgold, R. & Walker, M. P. The Neuroscience of Sleep. (Academic Press, 2010).

12. Crump, M. J. C. & Logan, G. D. Prevention and correction in post-error performance: An ounce of prevention, a pound of cure. Journal of Experimental Psychology: General 142, 692–709 (2013).

13. Laming, D. R. J. Information theory of choice-reaction times. (Academic Press, 1968).

14. Rabbitt, P. M. Errors and error correction in choice-response tasks. Journal of Experimental Psychology 71, 264–272 (1966).

15. Rabbitt, P. M. A. Error Correction Time without External Error Signals. Nature 212, 438–438 (1966).

16. Rabbitt, P. Psychological refractory delay and response-stimulus interval duration in serial, choice-response tasks. Acta Psychologica 30, 195–219 (1969).

17. Rabbitt, P. M. A. & Vyas, S. M. An elementary preliminary taxonomy for some errors in laboratory choice RT tasks. Acta Psychologica 33, 56–76 (1970).

18. Yamaguchi, M., Crump, M. J. C. & Logan, G. D. Speed-accuracy trade-off in skilled typewriting: decomposing the contributions of hierarchical control loops. J Exp Psychol Hum Percept Perform 39, 678–699 (2013).

19. Beilock, S. L., Carr, T. H., MacMahon, C. & Starkes, J. L. When paying attention becomes counterproductive: impact of divided versus skill-focused attention on novice and experienced performance of sensorimotor skills. J Exp Psychol Appl 8, 6–16 (2002).

20. Logan, G. D. & Zbrodoff, N. J. Stroop-type interference: Congruity effects in color naming with typewritten responses. Journal of Experimental Psychology: Human Perception and Performance 24, 978–992 (1998).

21. Logan, G. D. & Crump, M. J. C. The Left Hand Doesn’t Know What the Right Hand Is Doing: The Disruptive Effects of Attention to the Hands in Skilled Typewriting. Psychol Sci 20, 1296–1300 (2009).

22. Logan, G. D. & Crump, M. J. C. Cognitive Illusions of Authorship Reveal Hierarchical Error Detection in Skilled Typists. Science 330, 683–686 (2010).

23. Snyder, K. M. & Logan, G. D. The problem of serial order in skilled typing. Journal of Experimental Psychology: Human Perception and Performance 40, 1697–1717 (2014).

24. Snyder, K. M., Logan, G. D. & Yamaguchi, M. Watch what you type: The role of visual feedback from the screen and hands in skilled typewriting. Atten Percept Psychophys 77, 282–292 (2015).

25. van Beers, R. J., van der Meer, Y. & Veerman, R. M. What Autocorrelation Tells Us about Motor Variability: Insights from Dart Throwing. PLoS One 8, (2013).

26. Yamaguchi, M. & Logan, G. D. Pushing typists back on the learning curve: Contributions of multiple linguistic units in the acquisition of typing skill. Journal of Experimental Psychology: Learning, Memory, and Cognition 40, 1713–1732 (2014).

27. Oldfield, R. C. The assessment and analysis of handedness: The Edinburgh inventory. Neuropsychologia 9, 97–113 (1971).

28. Rachaveti, D. & Skm, V. Motor sequence learning data collected continuously for fifteen days of practice using a novel glove-based typing device. Data in Brief 29, 105234 (2020).

29. Fiehler, K., Ullsperger, M. & Cramon, D. Y. V. Electrophysiological correlates of error correction. Psychophysiology 42, 72–82 (2005).

30. Logan, G. D. & Crump, M. J. C. Chapter one - Hierarchical Control of Cognitive Processes: The Case for Skilled Typewriting. in Psychology of Learning and Motivation (ed. Ross, B. H.) vol. 54 1–27 (Academic Press, 2011).

31. Berniker, M. & Kording, K. Estimating the sources of motor errors for adaptation and generalization. Nature Neuroscience 11, 1454–1461 (2008).

32. Berniker, M. & Kording, K. P. Estimating the Relevance of World Disturbances to Explain Savings, Interference and Long-Term Motor Adaptation Effects. PLOS Computational Biology 7, e1002210 (2011).

33. van Beers, R. J. Motor learning is optimally tuned to the properties of motor noise. Neuron 63, 406–417 (2009).

34. Fitts, P. M. & Posner, M. I. Human performance. (Brooks/Cole, 1967).

35. Acuna, D. E. et al. Multifaceted aspects of chunking enable robust algorithms. J. Neurophysiol. 112, 1849–1856 (2014).

36. Newell, A. & Rosenbloom, P. S. The Soar Papers (Vol. 1). in (eds. Rosenbloom, P. S., Laird, J. E. & Newell, A.) 81–135 (MIT Press, 1993).

37. Logan, G. D. Toward an instance theory of automatization. in (1988). doi:10.1037/0033-295X.95.4.492.

38. Logan, G. D., Miller, A. E. & Strayer, D. L. Electrophysiological Evidence for Parallel Response Selection in Skilled Typists. Psychol Sci 22, 54–56 (2011).

39. Nissen, M. J. & Bullemer, P. Attentional requirements of learning: Evidence from performance measures. Cognitive Psychology 19, 1–32 (1987).

